## SupplementaryFigure1 for "Interactions between *Mycoplasma mycoides* subsp. *mycoides* and bovine macrophages under physiological conditions"

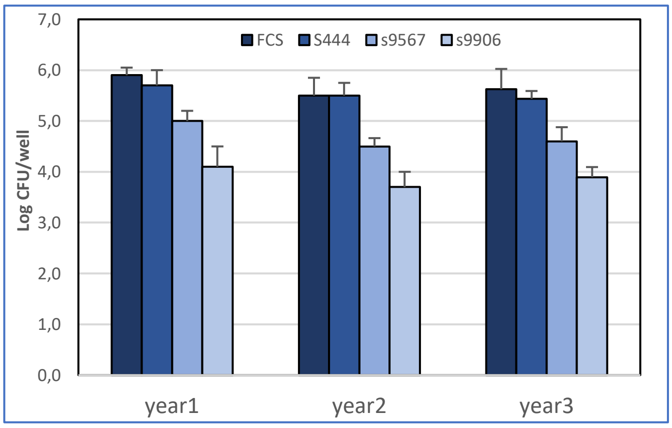


**Supplementary Figure 1**: Stability of the bactericidal effect of bovine complement. *Mmm* was incubated for 1h with decomplemented sera (FCS) or 10% of non-decomplemented bovine sera from 3 different animals (s+animal number). For each year, results from 3 different experiments are shown as mean+/-SD of CFU expressed in log10**.**
